# ATF5 is a regulator of exercise-induced mitochondrial quality control in skeletal muscle

**DOI:** 10.1101/2022.07.18.500448

**Authors:** Mikhaela B. Slavin, Rita Kumari, David A. Hood

## Abstract

**Objectives:** The Mitochondrial Unfolded Protein Response (UPR^mt^) is a compartment-specific mitochondrial quality control (MQC) mechanism that uses the transcription factor ATF5 to induce the expression of protective enzymes to restore mitochondrial function. Acute exercise is a stressor that has the potential to temporarily disrupt organellar protein homeostasis, however, the roles of ATF5 and the UPR^mt^ in maintaining basal mitochondrial content, function and exercise-induced MQC mechanisms in skeletal muscle are not known.

**Methods:** ATF5 KO and WT mice were examined at rest or after a bout of acute endurance exercise. We measured protein content in whole muscle, nuclear, cytosolic and mitochondrial fractions, in addition to mRNA transcript levels in whole muscle. Using isolated mitochondria, we quantified rates of oxygen consumption and ROS emission to observe the effects of the absence of ATF5 on organelle function.

**Results:** ATF5 KO mice exhibited a larger and less functional muscle mitochondrial pool, most likely a culmination of enhanced biogenesis via increased PGC-1α expression, and attenuated mitophagy. The absence of ATF5 resulted in a reduction in antioxidant proteins and increases in mitochondrial ROS emission, cytosolic cytochrome c, and the expression of mitochondrial chaperones. KO muscle also displayed enhanced exercise-induced stress kinase signaling, but a blunted mitophagic and UPR^mt^ gene expression response, complemented by significant increases in the basal mRNA abundance and nuclear localization of ATF4. Instead of promoting its nuclear translocation, acute exercise caused the enrichment of ATF5 in mitochondrial fractions. We also identified PGC-1α as an additional regulator of the basal expression of UPR^mt^ genes.

**Conclusion:** The transcription factor ATF5 retains a critical role in the maintenance of mitochondrial homeostasis and the appropriate response of muscle to acute exercise for the optimization of mitochondrial quality control.

**Graphical abstract:** A modified version of the schematic shown in Fig. 8 will be supplied at a later time.

## 1. Introduction

Skeletal muscle is an extremely malleable tissue and is able to alter its structural, physiological and metabolic phenotype in response to altered external demands. This “plasticity” of muscle [1] is due in part to the to the adaptability of the mitochondrial reticulum, being a large contributor to improvements in muscle endurance resulting from regular exercise [2,3]. Mitochondria not only produce cellular energy, but they also lie at the crossroads of a multitude of bidirectional signaling cascades with the nucleus. This “retrograde” (mitochondria-to-nuclear) communication induces specific transcriptional programs to monitor and preserve the organelle network [4,5], and serves as an important component of the mitochondrial quality control (MQC) systems. These include biogenesis (synthesis), mitophagy (degradation), antioxidant capacity and the maintenance of intra-organellar protein homeostasis [6,7].

Muscle contractions during voluntary exercise promote mitochondrial biogenesis and mitophagy, enhancing the quality of the mitochondrial pool [2,8–11]. At the onset of exercise, early signaling events are initiated, converging in part on the activation of the transcriptional co-activator PGC-1α [12–15]. This “master regulator” of organelle biogenesis coordinates the transcription of mitochondrial genes encoded by the nuclear genome (NuGEMPs), which require import into the mitochondria via specialized protein import machinery (PIM) components [16–18]. The mitochondrial proteome consists of ∼1200 proteins, and over 99% of these are transcribed in the nucleus, while less than 1% are encoded by mtDNA [19]. The regulation of these proteins, including their proper import, translation, folding, and degradation is referred to as protein homeostasis or “proteostasis” [20,21]. As an integral component of MQC, adequate proteostasis is critical for sustaining mitochondrial function, as dysregulation is linked to a variety of pathological conditions [22].

The status of the mitochondrial protein folding environment is reliant on the equilibrium between the abundance of internal misfolded proteins and the organelle’s intrinsic folding capacity. This is determined by the amount of chaperone and protease enzymes, which possess the dedicated functions of refolding and degrading misfolded proteins, respectively. Unaccustomed exercise may elicit a proteostatic imbalance, generated by an increase in misfolded proteins leading to acute organellar proteotoxic stress [23,24], ratifying a need for protein quality control strategies. As an adaptive mechanism, the cell activates a compartment-specific transcriptional program called the mitochondrial unfolded protein response (UPR^mt^) [25–27], named in relation to the UPR^ER^ [28]. Utilizing novel mitochondria-to-nuclear communication, the transcription of UPR^mt^ genes, including mitochondrial-specific chaperones and proteases, are increased to restore proteostasis within the organelle [29–31]. UPR^mt^ activation and the adequate expression of MQC enzymes are a critical adaptive mechanism for the maintenance of organelle quality and homeostasis [32–34].

Activating transcription factor 5 (ATF5) is a major regulator of the UPR^mt^ in mammalian cells, promoting the transcriptional induction of downstream UPR^mt^ targets during mitochondrial stress [35]. Basally, ATF5 is degraded in the mitochondrion, similar to ATFS-1 in *C. elegans* [28,29], while under stressful conditions, ATF5 exhibits retrograde translocation to the nucleus [35]. As part of the ATF/CREB family, ATF5 retains a DNA-binding domain allowing it to interact with the promoters of UPR^mt^ target genes, in addition to a bZIP domain facilitating its heterodimerization with other bZIP transcription factors [36,37]. Primarily abundant in the liver, ATF5 has been investigated in a plethora of cellular conditions. It is responsive during amino acid starvation [38–40], in mediating cell survival and proliferation in cancer and neurodegeneration [41–44], and in retaining an influential role in the maturation of olfactory sensory neurons (OSNs) [45]. However, evidence is beginning to emerge, supporting its role in mediating stress responses in muscle. ATF5 is upregulated in skeletal muscle during β-adrenergic stimulation [46], in mitochondrial myopathy [47], in myoblasts upon impaired protein translation [47], and during differentiation [48]. Furthermore, ATF5 has been found to be integral in UPR^mt^ activation and rescuing of mitochondrial function upon cardiac insult [49–51], providing opportunistic evidence for pursuing ATF5 function and UPR^mt^ activation in skeletal muscle.

The molecular mechanisms governing mitochondrial biogenesis in response to exercise are complex, and seminal work is beginning to reveal potential roles that acute mitochondrial proteotoxicity and the UPR^mt^ may have in mediating changes in cell signaling upon contractile activity [52–54]. Despite these findings, no work has focused on the role of ATF5 in the maintenance of basal mitochondrial homeostasis in muscle, or its function in mediating the UPR^mt^ in response to acute exercise. Therefore, we utilized WT and ATF5 KO mice to examine 1) the influence of ATF5 on mitochondrial content and function, 2) whether ATF5 is required for various MQC processes including biogenesis, mitophagy, antioxidant capacity, apoptosis, and 3) the role of ATF5 in exercise-induced UPR^mt^ activation and mitochondrial biogenesis signaling.

## 2. Materials and Methods

### 2.1. Animals

ATF5 whole-body KO mice were generated by crossing ATF5^tm1(KOMP)^ (Velocigene Project 11612) heterozygotes in a C57BL6/N background, generously provided by Dr. Stavros Lomvardas from Columbia University, with FVB WT females. Animals were housed in a 12:12-h light-dark cycle and given food and water ad libitum. To genotype progeny, ear clippings were obtained from each animal to make crude DNA extracts. They were subsequently mixed with JumpStart REDTaq polymerase (P0982, Sigma), forward and reverse primers (50 μM) (Table S1) for the WT and altered ATF5 gene, and subjected to amplification by PCR. Reaction products were run on a 1% agarose gel and visualized with the use of ethidium bromide. Experimental animals were used at 5.5-7 months of age and separated into either Control (CON) or Exercise (EX) groups (n=9-13/group). For the PGC-1α KO experiments, TA muscles were extracted from WT and whole-body PGC-1α KO mice at 4-5 months of age [55], snap-frozen in liquid nitrogen and stored at -80°C for later analysis by qPCR (n=3-5/group).

### 2.2. Exercise protocols

Two different acute exercise protocols were used in this study, where mice performed either a continuous, submaximal bout or an exhaustive exercise test. All exercised animals ran at a fixed 10% slope and were acclimatized to the treadmill for three days prior to exercising. The first was a bout of acute continuous exercise, where mice ran at a pace of 15 m/min for 60 minutes, followed by 18 m/min for 30 minutes (90 minutes total). The second was acute exhaustive exercise, which was completed as follows: 0 m/min for 5 minutes, 5 m/min for 5 minutes. 10 m/min for 5 minutes, 15 m/min for 5 minutes, 20 m/min for 5 minutes, 25 m/min for 5 minutes, increasing the speed by 1 m/min every 3 minutes until exhaustion. Exhaustion was defined as the animal remaining on the shock pad for 10 seconds, despite encouragement to run. Using a small tail bleed, blood lactate was measured with a Lactate Scout+ analyzer (EKF Diagnostics). The tissues were collected immediately after exercise and stored at - 80°C for later protein, mRNA, enzyme analyses, or used fresh for cellular fractionations.

### 2.3. Cytochrome c oxidase activity

Activity of the cytochrome c oxidase (COX) enzyme was used as a marker of mitochondrial content in muscle. Using a portion of the TA, tissues were placed in COX enzyme extraction buffer (100 mM Na-K-Phosphate, 2 mM EDTA, pH 7.2) on ice and diluted 40-fold. They were subsequently organized in metal brackets that were stored at -20°C and homogenized with stainless steel beads at 30 Hz using a TissueLyser II (Qiagen). Samples were lysed twice for one minute, followed by three to five 30-second rounds until fully homogenized. A test solution containing 20 mg of horse heart cytochrome c (C2506, Sigma) was prepared and incubated at 30°C. Using a multipipette, 240 μl of the test solution was added to 50 μl of whole muscle homogenate in a 96-well plate. The maximal oxidation rate of cytochrome c was then measured spectrophotometrically in a Synergy HT (Bio-Tek Instruments) plate reader, analyzing the change in absorbance at 550 nm and temperature of 30°C. For each sample, the COX activity measurement was determined as an average of three trials.

### 2.4. Nuclear and cytosolic fractionation

Nuclear and cytosolic fractions were prepared from one freshly extracted whole TA, using NE-PER Extraction Reagents (38835, Thermo Fisher Scientific) supplemented with phosphatase and protease inhibitors. Approximately ∼30-50 mg of tissue was minced on ice and homogenized in CER-I. Homogenates were then left to stand on ice for 10 minutes. Following the addition of CER-II, samples were briefly vortexed and centrifuged at 16,000 g for 10 minutes at 4°C. The supernatant (cytosolic fraction) was collected. The remaining pellets, containing nuclei and cellular debris, were washed 3 times in cold 1 x PBS and resuspended in NER. Nuclear fractions were then sonicated 3 times for 3 seconds each and incubated on ice for 40 minutes. Samples were vortexed every 10 minutes during the incubation, then underwent centrifugation at 16, 000 g for 10 minutes. The resulting supernates (nuclear fractions) were collected. Both cytosolic and nuclear subfractions were stored in -80°C until further analysis.

### 2.5. RNA isolation and reverse transcription

Approximately 50-70 mg of lysed gastrocnemius muscle tissue was combined with TRIzol® reagent (15596018, Life Technologies) and mixed with chloroform. Samples were centrifuged at 16,000 g for 15 minutes at 4°C, and the upper aqueous phase was transferred into a new tube with isopropanol and left overnight at -20°C to precipitate. Samples were once again centrifuged at 16,000 g for 10 minutes. The resulting supernatant fraction was discarded, and the pellet was suspended in 30 μl of molecular grade sterile H_2_O. RNA concentrations and purities were measured using the NanoDrop 2000 (Thermo Fisher Scientific). The Superscript III Reverse Transcriptase enzyme (Invitrogen) was used to reverse-transcribe 1.5 μg of RNA into cDNA.

### 2.6. mRNA expression using real-time PCR

The mRNA expression of *ATF5, ATF4, CHOP, PGC-1*α, *COX-IV, HSP60, LONP1, mtHSP70 and ClpP* was measured using the 7500 Real-Time PCR system (Applied Biosystems Inc.) and SYBR Green qPCR Master Mix (B21203, BiMake) in a 96-well plate. GAPDH and β-2 microglobulin were used as housekeeping genes for the normalization of transcript levels. Each well contained SYBR green, forward and reverse primers for the gene of interest (GOI) (20 μM) (Table 1), 10 ng of cDNA and sterile H_2_O to yield a final reaction volume of each well of 25 μl. Primer optimizations were run beforehand to control for primer dimers and nonspecific amplification by analyzing melt curves generated by the instrument. All samples were run in duplicates in tandem with negative control wells that contained sterile H_2_O in the place of cDNA.

### 2.7. Real-time PCR quantification

For the quantification of mRNA levels, the threshold cycle (C_T_) value of the GOI was first subtracted from the average C_T_ value of both endogenous reference genes to obtain the ΔC_T_ value for the GOI: ΔC_T_ = C_T_ (GOI) - C_T_ (reference). Next, the ΔC_T_ value of the exercised tissue (EX) was subtracted from the ΔC_T_ value of the control tissue (CON) for each genotype to obtain the ΔΔC_T_ values for the WT EX, KO CON, and KO EX samples: ΔΔC_T_ = ΔC_T_ (EX) - ΔC_T_ (CON). Results are expressed as fold changes above mRNA levels of WT CON animals using the ΔΔC_T_ method, calculated as 2^-^ΔΔ^CT^.

### 2.8. Immunoblotting

Whole muscle protein extracts prepared from a portion of the gastrocnemius and isolated subfractions were separated on 10-15% SDS-PAGE gels via electrophoresis at 120 volts for approximately ∼90 minutes, and subsequently transferred onto nitrocellulose membranes, stained with Ponceau Red and cut at the appropriate molecular weights. Blots were blocked in 5% skim milk or 5% BSA for phosphorylated proteins in TBS-T solution (25 mM Tris-HCl, 1 mM NaCl, 0.1% Tween-20, pH 7.5) for one hour at room temperature with gentle agitation. They were then coated with the appropriate primary antibodies (Table S2) and incubated overnight at 4°C. Membranes were then washed 3 × 5 minutes in TBS-T and incubated with HRP-linked secondary antibodies for one hour at room temperature. After another series of wash steps, blots were visualized with enhanced chemiluminescence using an iBright CL1500 Imaging System (Thermo Fisher Scientific). Quantifications were carried out using ImageJ (NIH) software and normalized to the corresponding loading controls or Ponceau. Corrected values were additionally normalized over a ‘Standard’ whole muscle sample that was run on every gel. For the cytochrome c blots, cytochrome c was probed for in cytosolic fractions isolated from the muscle of WT and ATF5 KO animals. Quantified values were corrected for VDAC and normalized for PFK-1. Approximate molecular weights in kDa (kilodaltons) of proteins are indicated on each blot.

### 2.9. Mitochondrial isolations

Approximately 700-1000 mg of fresh skeletal muscle tissue (one gastrocnemius, two quadriceps, and two triceps) was extracted from anesthetized animals, placed in ice-cold buffer, and subsequently minced and homogenized. Mitochondrial isolations were performed as previously described [56]. To separate SS and IMF mitochondria, homogenates were subjected to differential centrifugation at 800 g for 10 minutes. The IMF fraction was briefly treated with the Nagarse protease from *Bacillus lichenformis* (P5380, Sigma) to liberate the mitochondrial pool. After additional centrifugation steps, both SS and IMF pellets were suspended in resuspension buffer (100 mM KCl, 10 mM MOPS, 0.2% BSA pH of 7.4). Protein concentrations were determined using the Bradford method. Fresh mitochondrial fractions were utilized immediately to measure respiration and ROS emission.

### 2.10. Mitochondrial respiration

Oxygen consumption (O_2_/mg/min) over time in SS and IMF mitochondria was measured using a Clark Electrode (Strathkelvin Instruments). A small volume of either the SS or IMF fraction (50 μl) was incubated with 250 μl of VO_2_ buffer (250 mM sucrose, 50 mM KCl, 25 mM Tris base, 10 mM K_2_HPO_4_, pH 7.4) while being continuously stirred at 30°C. Respiration rates were determined (nanoatoms O_2_/min/mg) in the presence of 10 mM glutamate for State IV (passive) respiration, and 0.44 mM ADP for State III (active) respiration. Finally, NADH was added to test mitochondrial membrane integrity.

### 2.11. Mitochondrial ROS emission

SS and IMF mitochondria (75 μg) were incubated in a black polystyrene 96-well plate with VO_2_ buffer and 50 mM H_2_DCF-DA (D399, Thermo Fisher Scientific) at 37°C for 30 minutes. The fluorescence emission (480-520 nm) was measured in a Synergy HT (Biotek) plate reader using Gen5 software. ROS emission was assessed with the addition of 10 mM glutamate in the absence (State IV) or presence (State III) of 0.44 mM ADP. Data were normalized to the corresponding respiration rates.

### 2.12. Statistical analysis

Data were analyzed using GraphPad Prism 8.0 software and values are reported as means ±SEM. Basal comparisons of WT and KO animals were carried out with unpaired Student’s t-tests. A 2-way ANOVA was used in the comparisons of exercise conditions and genotypes, followed by a Bonferroni post-hoc test where necessary. Statistical significance was set at P<0.05.

## 3. Results

### 3.1. ATF5 KO mice do not exhibit differences in exercise tolerance and performance

The ATF5 KO mouse model was confirmed with DNA genotyping, through the presence of the mutant ATF5 gene and the absence of the WT gene (Fig. 1A) [50]. It was additionally confirmed with the abolishing of the ATF5 transcript in KO muscle (P<0.0001) (Fig. 1B). The phenotypic characteristics of ATF5 KO mice have been described previously [50]. In our study, deletion of the ATF5 gene resulted in a reduction in body weight of approximately 21% (P<0.0001) (Fig. 1C) without any changes occurring in relative muscle mass, observed by expressing TA weight per unit of body weight (Fig. 1D). To examine the involvement of ATF5 in determining exercise performance, WT and ATF5 KO mice were subjected to an incremental, exhaustive exercise bout. Animals were run to exhaustion, indicated by 3.8-4-fold increases in blood lactate post-exercise (P<0.0001) (Fig. 1E). No differences in exercise tolerance were observed between genotypes, measured through distance to exhaustion in the exhaustive exercise bout (Fig. 1F).

**Figure 1.**
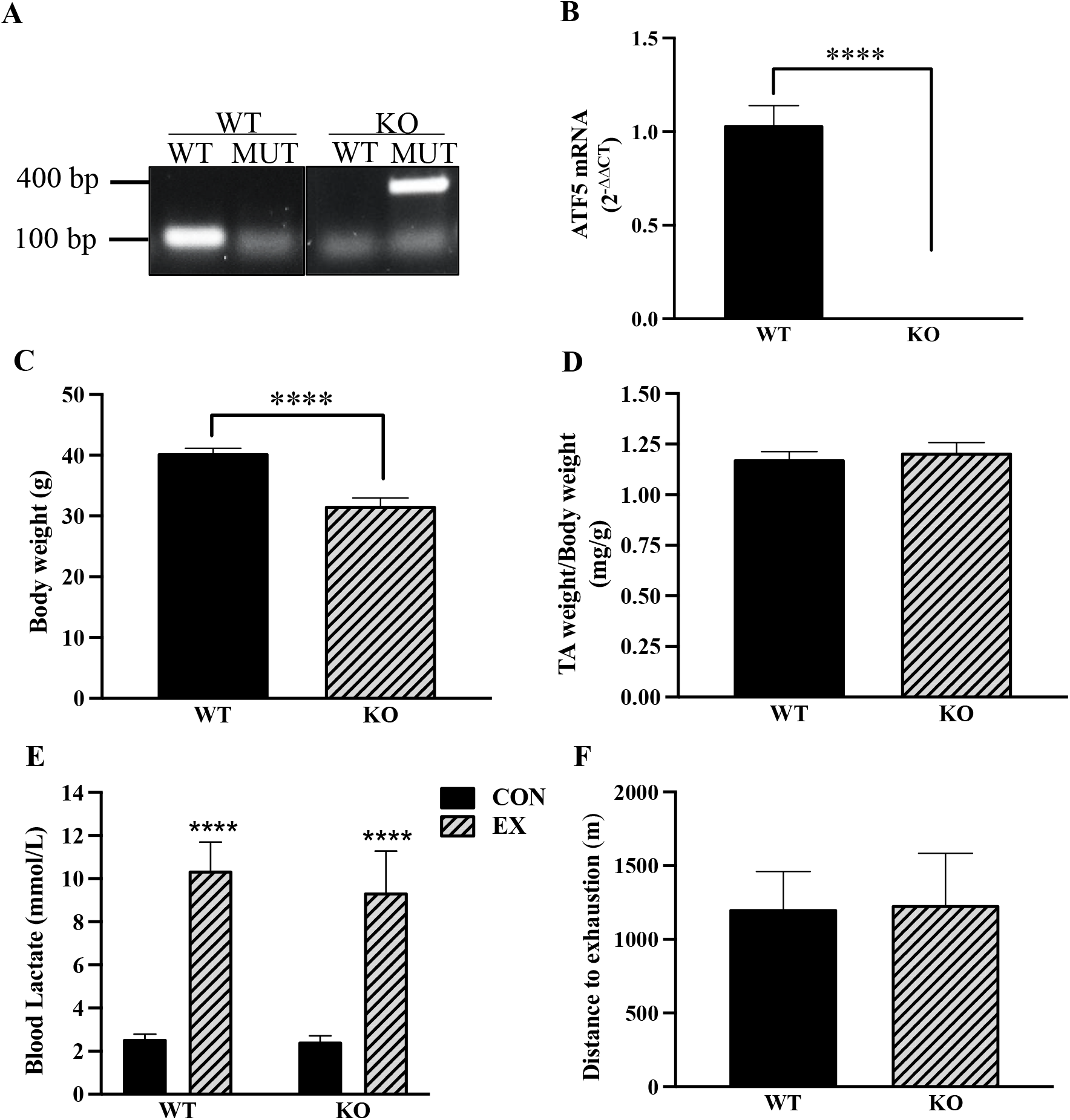
ATF5 KO animals exhibit reduced body weights with no changes in exercise tolerance or performance. The whole-body ATF5 KO mouse model was confirmed through **A)** DNA genotyping targeting the ATF5 WT (100 bp) and MUT (400 bp) alleles with designated primer sets and **B)** measuring ATF5 mRNA levels with qPCR. (WT: n=16, KO: n=23). Phenotypic characteristics of the genotypes were assessed, including **C)** Body weight and **D)** TA weight normalized for body weight, (WT: n=18, KO: n=26-27). **E)** Blood lactate levels in CON and EX animals subjected to acute exhaustive exercise (n=4-13). **F)** Exercise performance was observed from distance to exhaustion recordings from an incremental exhaustive exercise test on the treadmill (n=4-7). ****P<0.0001, unpaired t-test. bp, base pairs; CON, control; EX, exercised; KO, knockout; MUT, mutant; WT, wild-type.

### 3.2. asal protein expression of select UPR^mt^ chaperones is augmented with no changes in autophagy and lysosomal proteins in the absence of ATF5

Since it has been suggested that ATF5 is required for the expression of its downstream UPR^mt^ targets during mitochondrial stress [35], it was of interest to investigate the requirement of ATF5 in the basal expression of these proteins. Interestingly, expression of the UPR^mt^ chaperones HSP60 and Cpn10 was upregulated in ATF5 KO muscle by 1.4-fold (P<0.05) and 2.4-fold (P<0.05), respectively, along with a 1.8-fold increase in the UPR^mt^ transcription factor CHOP (P<0.05) (Figure 2A, C). However, no changes were observed in protein expression of the chaperone mtHSP70, the protease LONP1 or the transcription factor ATF4. Probing for autophagy and lysosomal markers to evaluate any basal differences in protein content between genotypes, we observed a 22% increase in ATG7 protein (P<0.05) in KO animals, with no changes in Beclin-1 or the lysosomal markers LAMP1 and V-ATPase (Fig. 2B,D).

**Figure 2.**
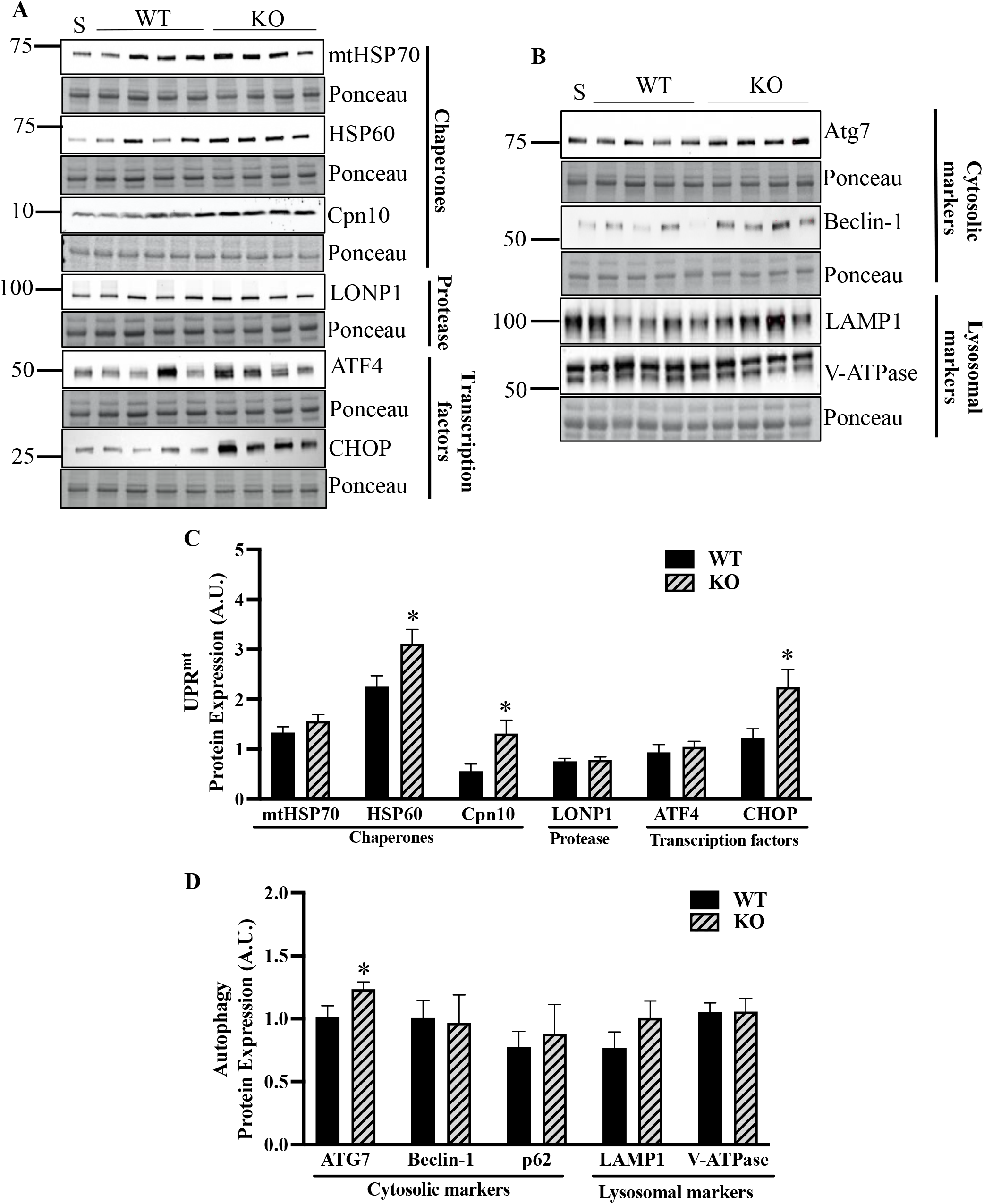
Increased protein expression of select UPR^mt^ markers in the absence of ATF5 with no changes in autophagy or lysosomal proteins in whole muscle. **A)** Representative blots of UPR^mt^ proteins in whole muscle with corresponding Ponceau stains. Proteins include chaperones mtHSP70, HSP60, Cpn10, the protease LONP1, and transcription factors ATF4 and CHOP. **B)** Representative blots of basal autophagy proteins in whole muscle, including cytosolic markers Atg7, Beclin-1 and lysosomal markers LAMP1 and V-ATPase with Ponceau stains. Corresponding quantifications of **C)** UPR^mt^ and **D)** autophagy and lysosomal proteins corrected for Ponceau (n=8-10). *P<0.05, unpaired t-test. A. U., arbitrary units; KO, knockout; S, standard; WT, wild-type.

### 3.3. The muscle of ATF5 KO animals displays enhanced mitochondrial content with reduced basal function

To analyze the influence of ATF5 on the maintenance of the mitochondrial pool, mitochondrial content was measured via protein content and enzyme activity analyses. In ATF5 KO muscle, there was a 43% increase in whole muscle PGC-1α protein (P<0.05) with no change in VDAC (Fig. 3A, B), indicating an increased drive for mitochondrial biogenesis. We also measured COX enzyme activity and performed mitochondrial yield calculations from mitochondrial isolations. KO muscle exhibited 14% greater COX activity relative to WT muscle (P<0.05) (Fig. 3C) and an increased mitochondrial yield of 24% (P<0.01) (Fig. 3D). We then investigated how the absence of ATF5 influences organelle function, as ATF5 is known to be required for the maintenance of basal mitochondrial function in HEK 293T cells in culture [35]. To investigate whether this holds true in muscle, oxygen consumption and ROS emission were measured in SS and IMF mitochondria isolated from the skeletal muscle of WT and ATF5 KO animals. In the absence of ATF5, significant reductions in mitochondrial respiration in State III conditions, but not State IV, were observed in both mitochondrial fractions derived from KO muscle. State III respiration was reduced by 40% in SS mitochondria (P<0.001) (Fig. 3E), and by 32% in the IMF pool (P<0.01) (Fig. 3F) in comparison to WT animals. Furthermore, SS mitochondria from KO muscle exhibited a trending increase in State IV ROS emission of 55% (P=0.06) (Fig. 3G) without any changes observed in IMF mitochondria (Fig. 3H).

**Figure 3.**
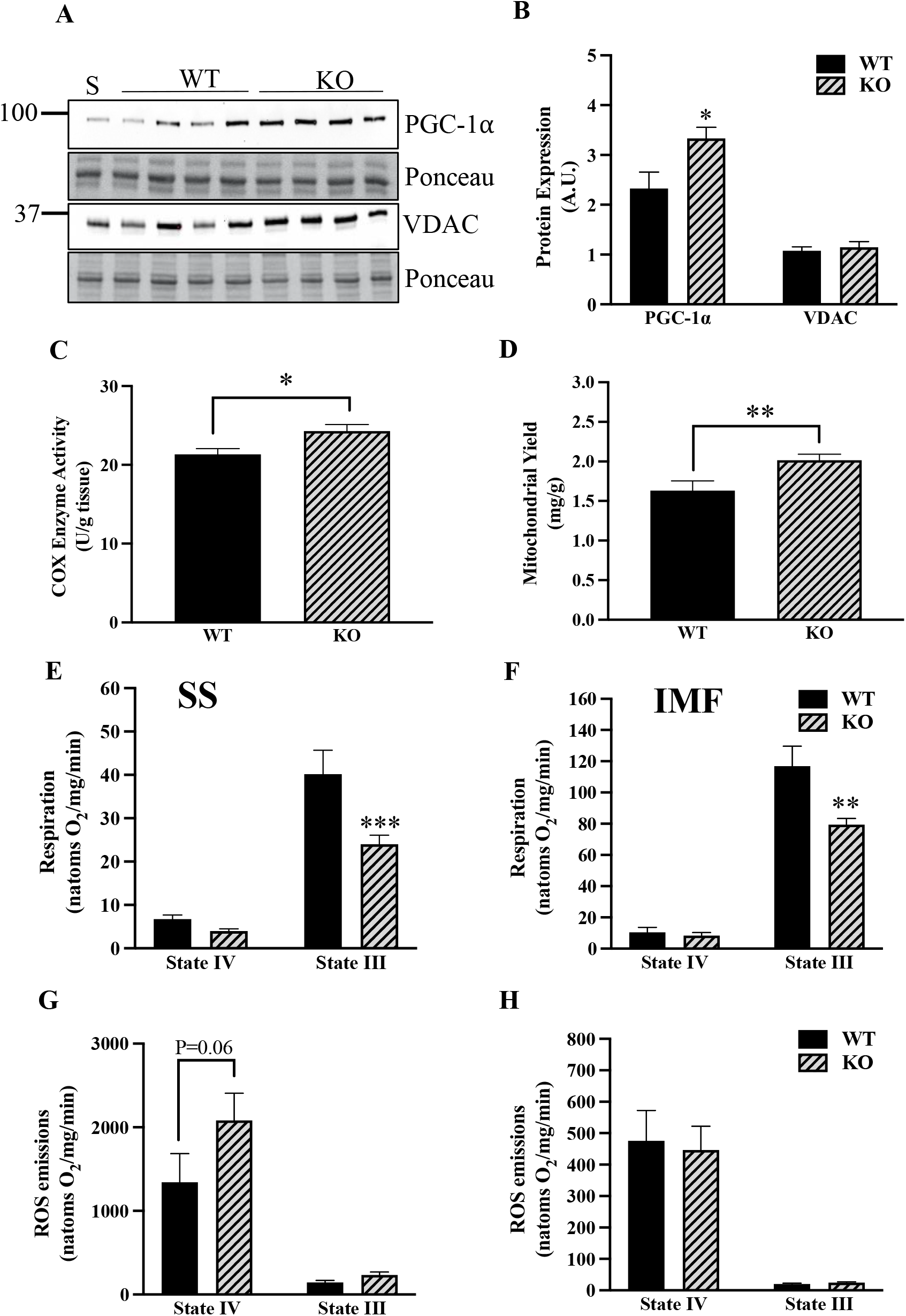
ATF5 KO muscle presents a more abundant mitochondrial pool with reduced organellar function. **A)** Representative blots in whole muscle of the transcription factor involved in mitochondrial biogenesis PGC-1α, the mitochondrial marker VDAC and corresponding Ponceau stains. **B)** Quantifications of PGC-1α and VDAC corrected for Ponceau (n=8-10). Mitochondrial content was assessed from **C)** COX Enzyme Activity values (n=9-10) and **D)** Mitochondrial Yields (SS and IMF combined) derived from mitochondrial isolation experiments (WT: n=18, KO: n=27). Mitochondrial respiration in **E)** SS and **F)** IMF mitochondria expressed in natoms O_2_/mg/min in both passive (State IV) and active (State III) respiratory conditions. Organelle function was also assessed by measuring ROS emission in **G)** SS and **H)** IMF subfractions (n=18-27). *P<0.05, **P<0.01, ***P<0.001 unpaired t-test, WT vs KO basally or in given respiratory state. A. U., arbitrary units; KO, knockout; S, standard; WT, wild-type.

### 3.4. The absence of ATF5 yields altered glycolytic enzyme levels, reduced antioxidant protein expression and increased cytochrome c release

ATF5 is known to have an impact on metabolism, as it regulates adipocyte differentiation and its knockdown in the white adipose tissue of mice improves whole-body insulin sensitivity [57]. To investigate how the absence of ATF5 regulates metabolic enzymes, the protein content of glycolytic enzymes GAPDH, PFK-1, LDHA and the β-oxidation enzyme CPT1b was measured in whole muscle protein extracts. ATF5 KO muscle exhibited a 27% increase in GAPDH (P<0.05) with a 14% reduction in the LDHA enzyme (P<0.05) (Fig. 4A, B), the LDH isoform responsible for the conversion of pyruvate to lactate. No changes in protein content were observed for PFK-1 or CPT1b. We also measured the levels of the cytosolic antioxidants NQO1, HO-1 and the mitochondrial antioxidant MnSOD in SS mitochondrial fractions. In whole muscle, NQO1 protein was reduced by 42% (P<0.05) in ATF5 KO samples with no changes in HO-1 (Fig. 4C, D). Furthermore, a reduction of 19% was observed in MnSOD in the SS pool derived from KO muscle (P<0.05) (Fig. 4C, D). The increases in mitochondrial ROS emission, coupled with these reductions in antioxidant enzymes in KO mice prompted us to question whether there was an increase in mitochondrially-mediated apoptotic signaling. Thus, we probed for cytochrome c in cytosolic samples fractionated from WT and ATF5 KO muscle (Fig. 4E, F). Interestingly, there was a 55% increase in cytosolic cytochrome c (P<0.05), potentially indicating increased mPTP opening and apoptotic signaling in the absence of ATF5.

**Figure 4.**
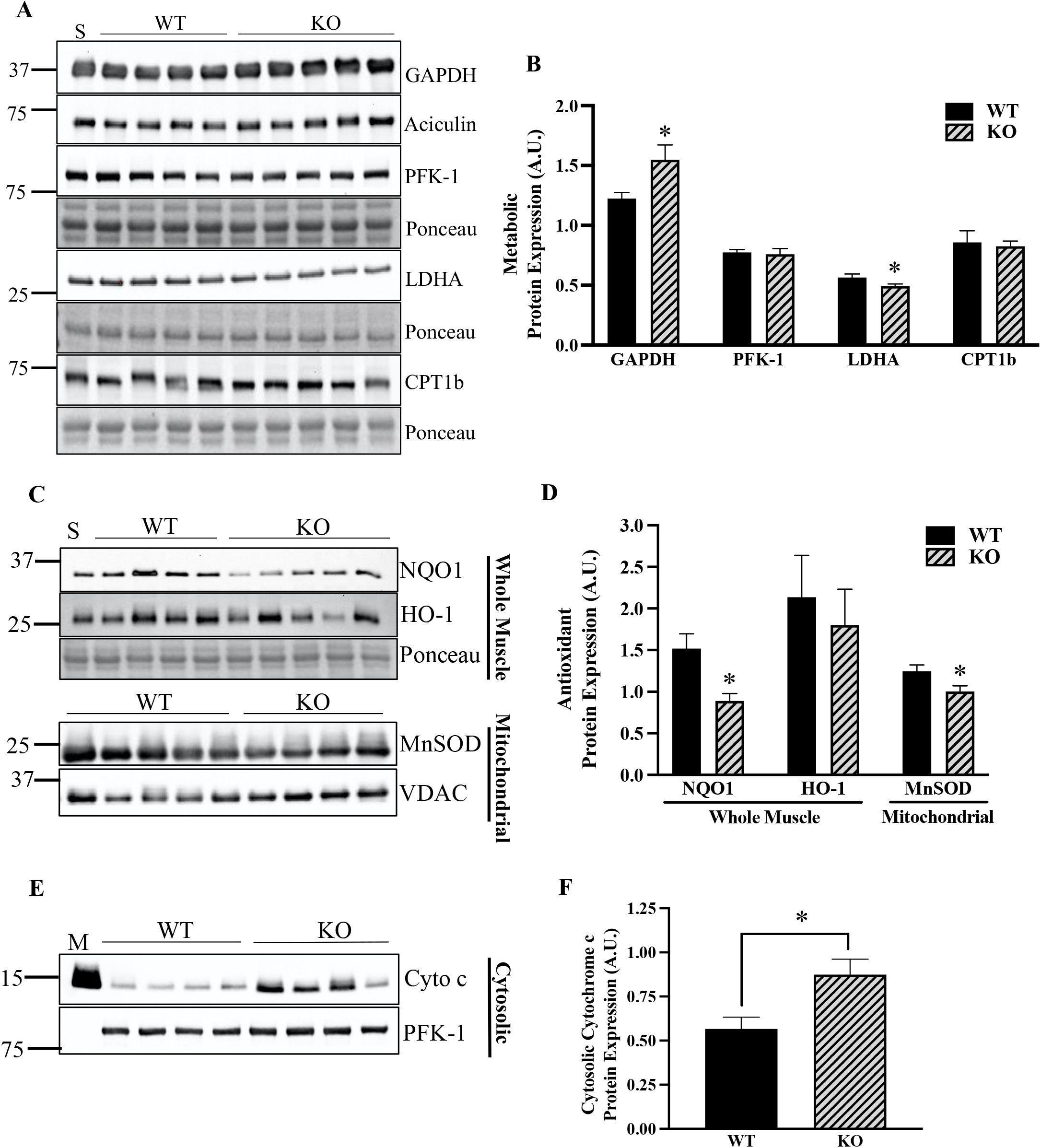
ATF5 regulates the expression of glycolytic and antioxidant enzymes, in addition to apoptotic cytochrome c release. **A)** Representative blots in whole muscle of the glycolytic enzymes GAPDH, PFK-1, LDHA and the β-oxidation protein CPT1b with Ponceau stains. **B)** Quantifications of the enzymes corrected for loading control or Ponceau (n=7-10). **C)** Representative blots of NQO1 and HO-1 in whole muscle samples with respective Ponceau stains, and MnSOD with VDAC in SS mitochondria. **D)** Quantifications of antioxidant proteins corrected for Ponceau stains or the respective loading control (n=6-9). **E)** Representative blot of cytochrome c and PFK-1 in cytosolic samples. **F)** Quantification of cytosolic cytochrome c protein corrected for PFK-1 (n=8). An amount of 5 μg was loaded to probe for GAPDH and was normalized to Aciculin, while 25 μg was used to target the other proteins and were normalized to Ponceau stains. *P<0.05 unpaired t-test. A.U., arbitrary units; M, IMF mitochondrial sample; KO, knockout; S, standard; WT, wild-type.

### 3.5. Acute exercise shifts ATF5 to the mitochondrial compartment, while ATF5 KO muscle exhibits blunted basal and exercise-induced mitophagy, and enhanced stress signaling with exercise

It has been previously documented in mammalian cells that various forms of mitochondrial stress induce the nuclear translocation of ATF5 [35]. Thus, we sought to investigate whether acute exercise was a sufficient stimulus to alter the cellular localization of ATF5 using western blotting in nuclear, cytosolic and mitochondrial fractions isolated from the muscle of control and exercised WT animals (Fig. 5A, B). Probing for ATF5 in nuclear and cytosolic fractions, there was no indication of its nuclear translocation. This prompted us to measure ATF5 levels in mitochondrial fractions, in which ATF5 protein content was enriched in the IMF fraction by 24% (P<0.05) (Fig. 5A, B).

**Figure 5.**
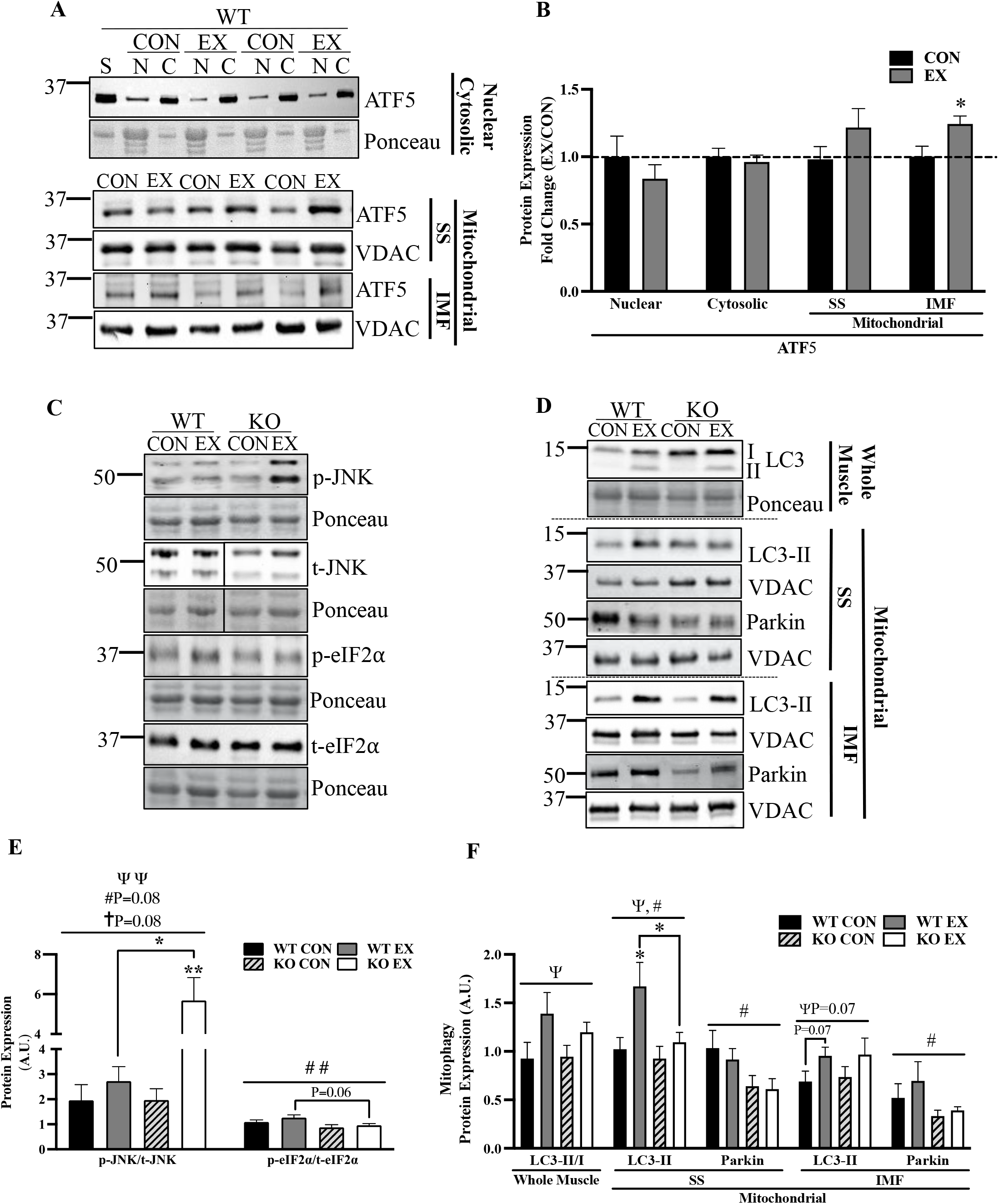
Acute exercise enhances mitochondrial ATF5 in WT mice, while there is reduced basal mitochondrial Parkin, attenuated exercise-induced mitochondrial LC3-II and enhanced stress signaling with exercise in ATF5 KO animals. **A)** Representative blots of ATF5 in nuclear, cytosolic and SS and IMF mitochondrial fractions from CON or EX animals subjected to acute exercise (continuous and exhaustive). The blot from nuclear and cytosolic samples is shown with the corresponding Ponceau stain, while that from mitochondrial samples is shown with VDAC. **B)** Graphical representation of ATF5 protein in different cellular fractions and conditions. Blots corrected for Ponceau or VDAC and displayed as a fold change (EX/CON) (n=6-9). Protein loaded for each fraction is as follows: Nuclear, 60 μg; Cytosolic, 10 μg; Mitochondrial, 25 μg. Representative blots of **C)** whole muscle p-JNK, p-eIF2α, t-JNK and t-eIF2α and **D)** autophagy proteins LC3-I, LC3-II and Parkin in whole muscle and SS and IMF mitochondrial samples in CON and EX animals subjected to acute exercise (continuous and exhaustive). Blots are shown with corresponding Ponceau stains or VDAC blots. **E)** Quantifications of phosphorylated/total for JNK and eIF2α corrected for Ponceau (n=5-12). **F)** Quantification of LC3 in whole muscle represented as the LC3-II/I ratio, and LC3-II and Parkin in SS and IMF mitochondria (n=6-11). ΨP<0.05, ΨΨP<0.01 main effect of exercise; #P<0.05, ##P<0.01 main effect of genotype; †P<0.05 interaction effect of exercise and genotype; *P<0.05, **P<0.01Bonferroni post-hoc analysis, CON vs EX of same genotype unless otherwise indicated. A. U., arbitrary units; C, cytosolic; CON, control; EX, exercised; KO, knockout; N, nuclear; p, phosphorylated; S, standard; t, total; WT, wild-type. Lines in blot indicate where two different areas were spliced together from the same blot.

The phosphorylation of the α subunit of ribosomal eIF2 at Serine 51 is a pivotal event during ISR activation, inhibiting global protein translation and selectively translating mRNAs with uORFs. These include the transcription factors ATF5, ATF4 and CHOP, which induce the transcription of mitochondrial-protective genes in the nucleus. On the other hand, JNK is a kinase that is highly responsive to acute exercise [58] and has been identified to be implicated in UPR^mt^ signaling during mitochondrial stress [59]. Thus, we sought to investigate whether there are discrepancies in acute exercise-induced UPR^mt^ upstream signaling, represented by eIF2α and JNK phosphorylation. While there was no significant effect of acute exercise on eIF2α phosphorylation, KO muscle had 29% less phosphorylated eIF2α relative to WT, with a significant effect of genotype across groups (P<0.01) (Fig. 5C, E). Exercise significantly increased JNK phosphorylation, and the magnitude of the effect appeared to depend on genotype (P=0.08), displaying a greater increase in KO animals. Post-hoc analyses showed that JNK phosphorylation increased by 2.9-fold (P<0.01) in KO mice, but only by 1.4-fold in WT muscle (Fig. 5C, E).

The response to acute exercise was further examined by investigating exercise-induced mitophagy. The more abundant mitochondrial pool in skeletal muscle of ATF5 KO mice could be a result of the increase in mitochondrial biogenesis indicated by more PGC-1α protein. However, we sought to investigate whether the increase in mitochondrial content could be a product of a reduction in mitochondrial degradation via mitophagy. Thus, we probed for LC3 in whole muscle samples and measured LC3-II and Parkin recruitment to mitochondria basally, and following acute exercise (Fig. 5D, F). There was a main effect of acute exercise in the LC3-II/LC3-I ratio in whole muscle (P<0.05), with increases of 50% and 27% in WT and ATF5 KO mice, respectively. To evaluate exercise-induced mitophagy, we probed for LC3-II in SS and IMF mitochondrial fractions. In SS mitochondria, there were main effects of exercise (P<0.05) and genotype (P<0.05) with a 64% increase in mitochondrial LC3-II in WT mice (P<0.05), but an increase of only 18% in KO animals. In the IMF fraction, there was a trending main effect of exercise (P=0.07) with a 39% increase in mitochondrial LC3-II in WT samples (P=0.07) and a similar 32% increase in KO muscle (Fig. 5D, F). Although acute exercise did not exert any changes in mitochondrial Parkin, there were significant effects of genotype in both SS and IMF mitochondria (P<0.05) (Fig. 5D, F). In KO muscle, basal mitochondrial Parkin was reduced by 33-38% and by 36-44% in SS and IMF fractions, respectively, when compared to WT muscle.

### 3.6. ATF5 KO muscle exhibits changes in the subcellular localization of PGC-1α, ATF4 and phosphorylated JNK2

In addition to whole muscle protein content analysis that was described above, the subcellular localization of PGC-1α ATF4, CHOP and p-JNK2 was measured in nuclear and cytosolic fractions isolated from WT and KO muscle following the exhaustive exercise test. A main effect of genotype was observed for both cellular compartments for PGC-1α protein. Nuclear PGC-1α was enhanced by 2.1-fold basally in the KO animals in comparison to WT counterparts, (P<0.05) (Fig. 6A, B), while collectively, cytosolic PGC-1α was increased by 49% when combining values from both control and exercised animals (P<0.05) (Fig. 6A, C). However, proportions of nuclear PGC-1α relative to total PGC-1α did not differ significantly between genotypes (Fig. 6D). With regards to the cellular localization of ATF4, 10-14% of total ATF4 was present in the nucleus in WT muscle, while KO muscle exhibited its enhanced nuclear translocation, with its nuclear proportions being 19-24% (P<0.05) (Fig. 6D). We did not observe any influence of acute exercise on the cellular localization of ATF4. However, in WT animals, nuclear CHOP increased by 59% following acute exercise (P<0.05) but only by 13% in KO mice. We also investigated the cellular localization of phosphorylated JNK2, as total JNK1 did not appear in nuclear fractions (Fig. 6A). Approximately 55-60% of p-JNK2 in the cell was found in the nucleus (Fig. 6D). Although there were no effects of exercise, collectively, the KO animals exhibited 74% more nuclear p-JNK2 than the WT mice (P<0.05) (Fig. 6B).

**Figure 6.**
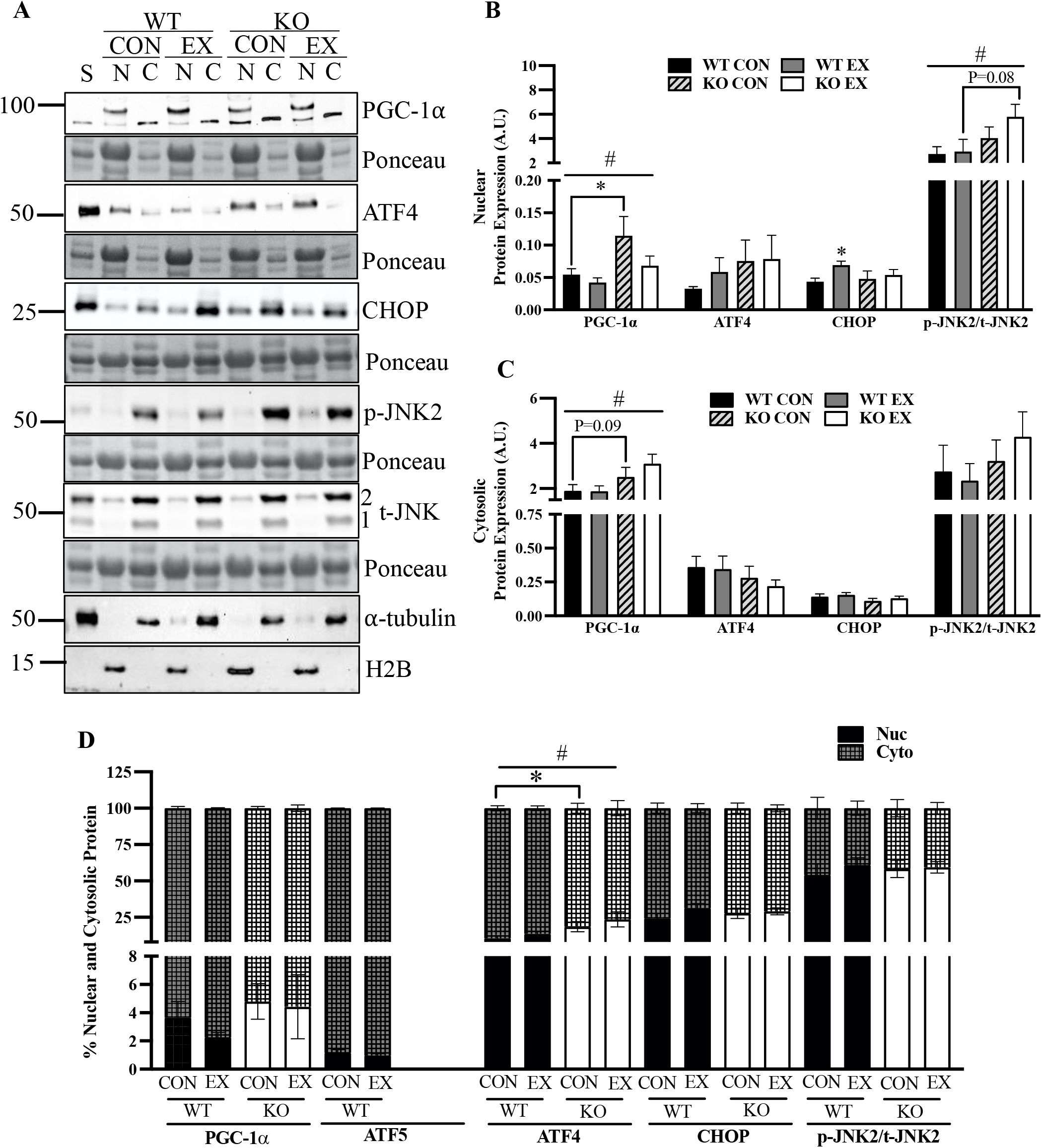
A lack of ATF5 in muscle elicits alterations in the cellular localization of proteins. **A)** Representative blots of PGC-1α, ATF4, CHOP, p-JNK2 (∼54 kDa) and t-JNK2 (∼54 kDa) in nuclear and cytosolic fractions of CON and EX animals subjected to acute exhaustive exercise. Blots are shown with corresponding Ponceau stains and blots of α-tubulin and H2B are also shown as indicators of sample purity. Quantifications shown for **B)** Nuclear and **C)** Cytosolic protein corrected for Ponceau stains. **D)** Cellular translocation data expressed as a percentage of protein in either cellular compartment (n=4-9). #P<0.05 main effect of genotype; *P<0.05 Bonferroni post-hoc analysis, CON vs EX of same genotype unless otherwise indicated. A.U., arbitrary units; C, cytosolic; CON, control; EX, exercised; KO, knockout; N, nuclear; p, phosphorylated; S, standard; t, total; WT, wild-type.

### 3.7. ATF5 KO animals exhibit attenuated exercise-induced UPR^mt^ signaling, while PGC-1α KO mice display reduced basal expression of UPR^mt^ genes

In mammalian cells in culture and in cardiac injury in rodents, ATF5 is known to be required for the stress-induced transcriptional activation of its downstream targets, including HSP60, LONP1 and mtHSP70 [35,50]. Thus, we investigated whether ATF5 KO animals exhibit impaired transcriptional signaling through the measurement of mRNA levels in control animals, as well as those subjected to a bout of continuous endurance exercise, with tissues collected immediately post-exercise (Fig. 7A). As an indicator of mitochondrial biogenesis signaling, there was a main effect of exercise (P<0.01) as PGC-1α mRNA was enhanced in both animal models after acute exercise, increasing by 5.8-fold (P<0.05) and 3.1-fold (P<0.05) in WT and KO animals, respectively (Fig. 7A). A main effect of exercise was also observed for COX-IV mRNA (P<0.05), as a 40% reduction was observed in KO muscle post-exercise (P<0.05) with no changes occurring in WT animals (Fig. 7A).

**Figure 7.**
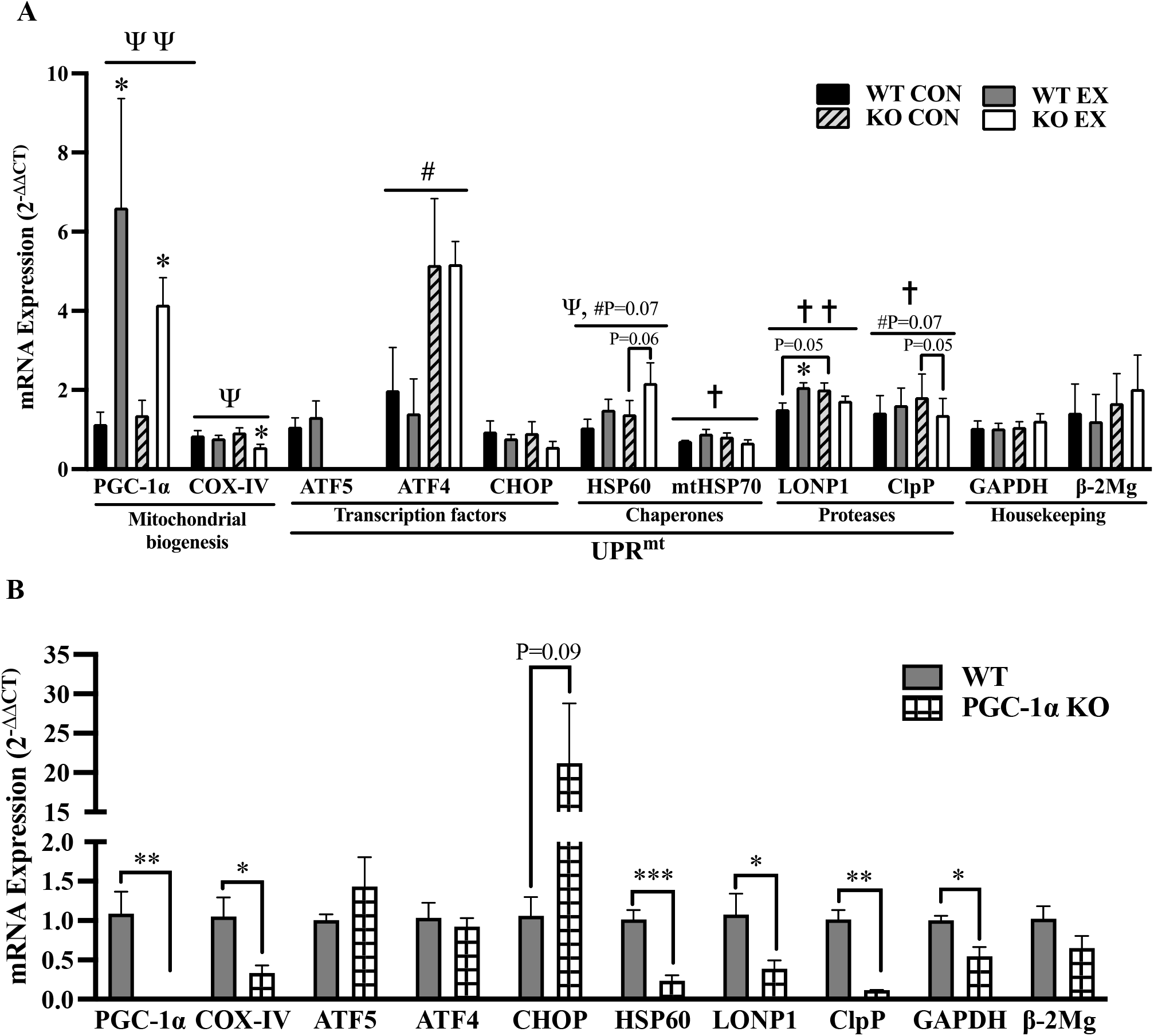
ATF5 is required for the increase in UPR^mt^ mRNAs with acute exercise, while PGC-1α coordinates their basal expression. **A)** Quantification of mRNA levels of genes involved in the mitochondrial gene expression response to acute continuous exercise. Targeted genes include mitochondrial biogenesis markers PGC-1α, COX-IV; UPR^mt^ transcription factors ATF5, ATF4, CHOP; UPR^mt^ chaperones HSP60, mtHSP70; UPR^mt^ proteases LONP1 and ClpP (n=4-5). ΨP<0.05, ΨΨP<0.01 main effect of exercise; #P<0.05 main effect of genotype; †P<0.05, ††P<0.01 interaction effect of exercise and genotype; *P<0.05 Bonferroni post-hoc analysis, CON vs EX of same genotype unless otherwise indicated. **B)** Quantification of basal mRNA levels of mitochondrial and UPR^mt^ genes in WT and PGC-1α KO muscle. PGC-1α, COX-IV, ATF5, ATF4, CHOP, HSP60, LONP1 and ClpP (n=3-5). *P<0.05, unpaired t-test. GAPDH and β2-Microglobulin served as housekeeping genes in both experiments. Values are expressed using the 2^-ΔΔCT^ method. CON, control; EX, exercised; KO, knockout; WT, wild-type.

No significant changes were observed in ATF5 or ATF4 mRNA with exercise. However, ATF4 mRNA was 2.6-fold and 3.7-fold higher in KO control and exercised samples, respectively, relative to WT mice (P<0.05). No changes in CHOP mRNA were observed between WT and KO mice basally, or with exercise. There was a main effect of exercise for HSP60 mRNA (P<0.05) as it was increased by 43% and 57% in WT and KO mice, respectively, and was more apparent in the KO animals (P=0.06). In addition, there was a trending main effect of genotype (P=0.07), with 32% and 45% more HSP60 mRNA in control and exercised KO animals, respectively.

The gene expression response of UPR^mt^ markers downstream of ATF5 other than HSP60, such as mtHSP70, LONP1 and ClpP, was also evaluated in WT and KO animals. Basal mRNA levels of LONP1 and ClpP were elevated by 33% (P=0.05) and 27% (P=0.07) in KO muscle, respectively. In contrast to what was observed with HSP60, the absence of ATF5 in muscle had a divergent response of these UPR^mt^ mRNAs to exercise, culminating in significant interaction effects of exercise and genotype for mtHSP70 (P<0.05), LONP1 (P<0.01) and ClpP (P<0.05). These targets increased by 13-37% in WT animals but were reduced by 14-25% in KO mice.

As the “master regulator” of mitochondrial biogenesis, PGC-1α coordinates the expression of many mitochondrial genes, and has also been shown to mediate increases in genes of the UPR^ER^ with acute exercise in mouse muscle [53]. Since PGC-1α protein was upregulated in both nuclear and cytosolic compartments in ATF5 KO animals, we investigated whether PGC-1α coordinates the expression of UPR^mt^ genes by measuring mRNA levels of target genes in WT and PGC-1α KO muscle tissues (Fig. 7B). PGC-1α mRNA was absent in PGC-1α KO mice as expected (P<0.01), while that of its bona fide downstream target, COX-IV of the electron transport chain, was reduced by 68% (P<0.05). Although no significant changes were observed in the expression of the transcription factors ATF5 or ATF4, CHOP appeared to be enhanced by 20-fold in PGC-1α KO muscle (P=0.09). The downstream UPR^mt^ targets HSP60, LONP1 and ClpP were significantly downregulated in the absence of PGC-1α, by 77% (P<0.001), 64% (P<0.05) and 89% (P<0.01), respectively.

## 4. Discussion

The maintenance of mitochondrial quality via the balance of biogenesis and mitophagy pathways in muscle has received considerable attention in recent years. In contrast, investigations into the contribution of protein quality control mechanisms to the maintenance of organelle function in mammalian organ systems have only recently begun to emerge, specifically with regard to the UPR^mt^. ATF5 is a transcription factor that has been found to be a mediator of the UPR^mt^ by translocating to the nucleus during various forms of mitochondrial stress to stimulate the transcription of chaperones and proteases [35]. Recent findings have emphasized the importance of an ATF5-dependent UPR^mt^ in rescuing cardiac function during injury [50,51]. In light of this, we questioned whether the requirement of this mechanism is conserved in skeletal muscle during exercise stress. Our previous research in this area has shown that an induction of UPR^mt^ occurs in response to muscle contractile activity, and that this actually precedes signaling towards mitochondrial biogenesis in a rodent model [52]. This suggests the importance of upregulating the expression of proteostatic machinery, possibly via the action of ATF5, for mitochondrial adaptations to exercise to occur. Thus, using ATF5 KO mice, we evaluated basal and exercise-induced mitochondrial quality control processes. Our results indicate that ATF5 is required for the maintenance of basal mitochondrial function, as well as exercise-induced UPR^mt^ mRNA and mitophagic response. Furthermore, ATF5 may have a regulatory role in substrate metabolism, antioxidant capacity and apoptosis, revealing ATF5 to have an influential role in muscle physiology (Fig. 8).

**Figure 8.**
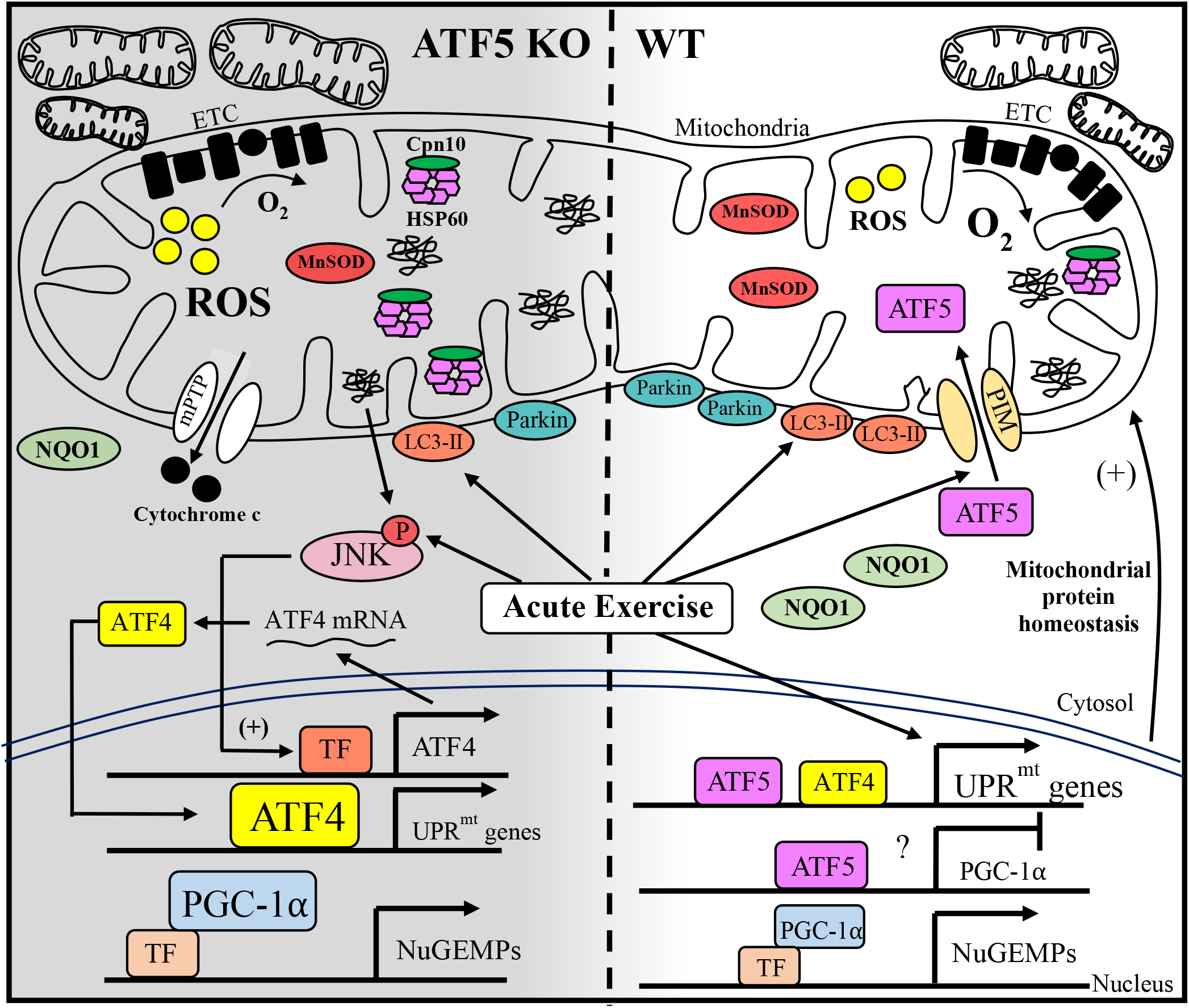
Working model illustrating the altered physiology of ATF5 KO muscle. The absence of ATF5 in skeletal muscle yields a more abundant mitochondrial pool composed of organelles that are less functional. This is characterized by reductions in oxygen consumption and enhanced ROS emission in comparison to WT mitochondria. Increases in ROS may be exacerbated by decreases in the expression of the antioxidant enzymes MnSOD and NQO1, inducing an increase in apoptotic cytochrome c release into the cytosol via the mPTP. The increased abundance of nuclear PGC-1α in the absence of ATF5 may be contributing to the transcription of nuclear genes encoding mitochondrial proteins (NuGEMPs) as well as increases in mitochondrial content observed, suggesting that ATF5 may be a negative regulator of PGC-1α in WT conditions. An enlarged mitochondrial pool in KO muscle may also be attributed to decrements in basal mitophagy indicated by reduced mitochondrial Parkin. ATF5 KO animals also exhibit a blunted mitochondrial quality control response to acute exercise stress, with an attenuated induction of mitochondrial LC3-II and the transcription of UPR^mt^ mRNAs. However, the increased mRNA levels and nuclear localization of ATF4 may explain the enhanced expression of chaperones HSP60 and Cpn10 basally in ATF5 KO muscle. Despite the attenuated mitochondrial response (above), an increased stress kinase signaling was evident post-exercise in ATF5 KO muscle, represented by enhanced JNK phosphorylation. The activation of JNK could also result from mitochondrial proteotoxicity to induce the transcription of ATF4 and the increase in ATF4 expression. Finally, acute exercise appears to induce the import of ATF5 into mitochondria, rather than to the nucleus. Solid arrows indicate evidence that signaling is occurring.

Given the function of ATF5 in regulating the UPR^mt^, we originally hypothesized that the absence of this transcription factor would result in a decrease in mitochondrial content. However, our findings indicate that the muscle of ATF5 KO animals displayed increased mitochondrial content in comparison to WT animals. We then assessed whether these mitochondria were equally functional, by measuring respiration and ROS emission in isolated mitochondria. Our data show that mitochondria from ATF5 KO muscle displayed reduced rates of oxygen consumption, and increases in ROS emission, most likely a product of reduced antioxidant capacity, in comparison to mitochondria from WT animals. These findings support other work that measured mitochondrial respiration in the absence of ATF5 in a mammalian cell model [35], and they indicate that in the absence of ATF5, skeletal muscle possesses a larger mitochondrial pool comprised of more dysfunctional mitochondria. The result of this compensatory adaptation is little to no change in exercise capacity between the two genotypes. Whether this altered mitochondrial content and function is confounded by inherent differences in the habitual activity or inactivity level of the KO animals remains to be determined.

To further investigate the underlying reasons for this increase in organelle content, we examined the expression and localization of PGC-1α. We found increased expression in whole muscle samples, enhanced in both the nuclear and cytosolic fractions isolated from ATF5 KO muscle, relative to WT samples. Thus, an increased transcriptional drive for mitochondrial biogenesis in the absence of ATF5 is contributing to a larger mitochondrial pool, albeit composed of more dysfunctional organelles. This indicates that ATF5 may be a negative regulator of PGC-1α and oxidative genes, similar to what has been observed with respect to its fellow bZIP transcription factor ATF4, which has recently been shown to be a negative regulator of Tfam and consequently, the expression of mtDNA-derived proteins [32]. Supporting this, ATFS-1, the nematode homologue of ATF5, is known to negatively regulate the expression of OXPHOS genes encoded by both nuclear and mitochondrial genomes [60].

An accumulation of poor quality mitochondria in the absence of ATF5 may also be a consequence of impaired mitophagy, the process whereby defective mitochondria are degraded and processed by the lysosomes [61]. The involvement of ATF5 in mitophagy is suggested by the fact that ATF5 levels are increased when mitophagy is inhibited in the myocardium, along with other UPR^mt^ proteins, possibly playing a compensatory role in the maintenance of mitochondrial homeostasis during cardiac stress [62]. Both SS and IMF mitochondrial subfractions isolated from ATF5 KO muscle display reduced levels of Parkin, an E3 ubiquitin ligase involved in mitophagy [9]. Based on this, organelles in ATF5 KO muscle may be less equipped to undergo basal mitophagic degradation, contributing to the enlarged pool of less functional mitochondria. Acute exercise also stimulates the recruitment of LC3-II to induce mitophagy to clear out dysfunctional organellar components of the muscle pool [15,63]. The SS fraction isolated from ATF5 KO muscle displayed attenuated recruitment of LC3-II following acute exercise, a similar phenomenon that we have observed in other animal models [9,15] corresponding with a reduction in mitophagy flux. Thus, in the absence of ATF5, mitochondria are less primed to be cleared via mitophagy to maintain the normal and exercise-induced regulation of mitochondrial quality control. The absence of changes in lysosomal proteins in ATF5 KO muscle suggests that ATF5 influences upstream autophagic signaling, perhaps with respect to autophagosomal formation, instead of the degradative end-stages. In future studies it will be interesting to see if the endurance training-induced increase in mitochondrially-localized Parkin that we have previously observed [9] can rescue the apparent stress-induced mitophagy defect observed here.

Among the many cellular events that encompass the physiological response to the stress of acute exercise, the activation (phosphorylation) of stress-inducible kinases is at the forefront of this molecular cascade. Our data show that acute exercise increases JNK activation, a member of the MAPK family of kinases, that has been shown to contribute to increases in PGC-1α gene expression following exercise [12,64,65]. We detected two isoforms of JNK in muscle, JNK1 (∼46 kDa) and JNK2 (∼54 kDa) with similar but yet distinct functions in some signaling pathways [66]. The JNK activation response to acute exercise is exaggerated in ATF5 KO muscle, in addition to its enhanced nuclear abundance. Interestingly, we did not detect JNK1 in the nucleus, and thus, only quantified nuclear JNK2. Regardless, JNK2 is the isoform that is implicated in UPR^mt^ signaling and it is responsive to mitochondrial proteotoxicity, lying downstream of the ISR kinase, PKR [59]. JNK activation is also highly responsive to ROS production [67,68], and could be a consequence of the elevated ROS levels that we observed in SS mitochondria from ATF5 KO muscle. We have shown that enhancing ROS levels induces increases in mitochondrial cytochrome c release and JNK phosphorylation in muscle to induce apoptosis [68]. Thus, the elevations in ROS emission and increased JNK signaling could explain the greater release of cytochrome c in ATF5 KO mice, culminating in its higher abundance in the cytosol. This could indicate an increase in apoptotic activation in ATF5 KO muscle, coinciding with the anti-apoptotic function of ATF5 that has been observed in other studies. [43,69].

In contrast to our initial hypothesis, protein markers previously established to be downstream targets of ATF5, such as mitochondrial chaperones and proteases [35], were increased basally in ATF5 KO muscle. To investigate the reason for this, we studied the expression and localization of ATF4, a well-established transcription factor that regulates the UPR^ER^ [70]. Sharing close homology with ATF5, ATF4 controls the transcription of ATF5, is activated by endoplasmic reticulum (ER) and mitochondrial stress [71,72], and its protein levels are upregulated in mammalian cells in the absence of ATF5 [73]. Whether ATF4 truly regulates the transcription of chaperones and proteases of the UPR^mt^ remains controversial [71,72], however our results indicate that muscle from ATF5 KO animals exhibited enhanced ATF4 mRNA levels, in addition to its increased nuclear localization. These changes suggest that either ATF5 negatively regulates the transcription of ATF4, or that signals consequent to the lack of ATF5 promote ATF4 expression and localization.

The ATF5 protein harbours both a mitochondrial targeting and nuclear localization sequence that regulates its organelle-partitioning depending on the cellular environment, as does its worm homologue ATFS-1 [74]. A previous study showed that cellular proteotoxicity induces the nuclear translocation of ATF5 due to its failed mitochondrial import [35]. However, acute exercise failed to induce this, and instead increased the abundance of ATF5 in IMF mitochondria. Thus, it is possible that a greater accumulation of misfolded proteins is required to inhibit the mitochondrial import of ATF5 and prompt its nuclear translocation immediately following exercise stress. This is a similar phenomenon to what has been observed in a yeast model, identifying an “early” and “late” UPR^mt^ with diverging efficiencies of mitochondrial protein import depending on the amount of time elapsed post-stress, as well as and the intensity of the stimulus imposed [75]. It is also known that chronically-imposed exercise induces an increase in mitochondrial protein import [76,77]. Whether the import flux is regulated by acute exercise to facilitate ATF5 accumulation inside the organelle remains a compelling question.

Previous research has identified that the expression of UPR^mt^ markers, including chaperones and proteases, is extremely responsive to inducible mitochondrial stressors, in an ATF5-dependent manner [35]. Thus, we sought to investigate whether acute exercise influences UPR^mt^ signaling skeletal muscle, and whether ATF5 is required for this response. The exercise protocol chosen was a sufficient exercise stimulus, as it augmented the transcript levels of PGC-1α in both genotypes, a shown previously [15,55]. Our data also indicate that ATF5 is not required for the induction of PGC-1α signaling following acute exercise, confirming what has also been shown in cardiomyocytes during cardiac stress [49].

However, WT and ATF5 KO mice displayed markedly divergent exercise-induced changes in mRNA transcripts of the UPR^mt^ downstream markers mtHSP70, LONP1 and ClpP. The upregulation in UPR^mt^ transcripts in WT muscle suggests that acute exercise is a sufficient stimulus to improve the protein folding capacity of mitochondria, similar to what occurs with respect to mRNA expression of UPR^ER^ markers [53]. Despite the relatively low abundance of ATF5 detected in the nuclear fractions, these levels are clearly sufficient to induce this exercise stress response in WT animals, which is absent in KO muscle. These results coincide with the concept that ATF5 may be part of a team of transcription factors that regulates the basal expression of UPR^mt^ markers [35], but that it is required for normal stress-induced changes in these downstream proteins. Other studies have confirmed that ATF5 is dispensable for the expression of downstream targets under basal conditions, but that it is required during a plethora of mitochondrial stressors, including proteotoxicity and ROS, to activate the UPR^mt^ and rescue mitochondrial function in mammalian cells and the mouse heart [35,50]. Interestingly, digressing from this collective response is the change in HSP60 gene expression, an established UPR^mt^ chaperone which is extremely responsive to acute exercise. However, these changes occurred in WT animals and were even more pronounced in the absence of ATF5, suggesting redundancies in the transcriptional regulation of HSP60 during exercise stress that do not obligate ATF5. This alternative regulation of HSP60 in the absence of ATF5 in comparison to the other proteins is also supported by the enhanced basal expression of HSP60 protein and that of its co-chaperone, Cpn10. Our data also reveal that HSP60 mRNA and that of other downstream UPR^mt^ markers were severely downregulated in PGC-1α KO muscle. Since PGC-1α is upregulated in ATF5 KO tissue, PGC-1α could be driving the increase in HSP60 mRNA and protein, with a heavier reliance of other UPR^mt^ genes on the presence of ATF5 for their transcription during exercise stress. PGC-1α has been shown to be required for the induction of UPR^ER^ genes post-exercise, but is not required for their basal expression. Thus, the novelty of our data supports PGC-1α as having a compartment-specific role within the cell to regulate the expression of protein folding under basal physiological conditions.

### 5.0. Conclusion

Taken together, our results indicate that the skeletal muscle of ATF5 KO animals exhibits 1) enhanced acute exercise-induced stress signaling, 2) increased basal expression of UPR^mt^ proteins, PGC-1α and ATF4, 3) a more abundant mitochondrial pool, with reduced organelle function, and 4) an altered UPR^mt^ gene expression and mitophagy response to acute exercise. This emphasizes ATF5 to be a critical regulator of mitochondrial quality control in skeletal muscle. In future investigations, it is worth pursuing the question of whether ATF5 is required to mediate beneficial mitochondrial adaptations to endurance training. Finally, it will also be compelling to examine the importance of ATF5 expression in coordinating the action of the UPR^mt^ upon impaired mitochondrial protein import and induced proteotoxicity, to preserve organelle function in skeletal muscle.

## Abbreviations

ANOVA: Analysis of variance
ARE: ATF5-specific response element
ATF: Activating transcription factor
ATFS-1: Activating transcription factor associated with stress-1
ATG7: Autophagy-related protein 7
bZIP: Basic leucine zipper
C/EBP: CCAAT-enhancer binding protein
CCA: Chronic contractile activity
cDNA: Complimentary DNA
CHOP: C/EBP homologous protein
ClpP: Caseinolytic mitochondrial matrix peptidase proteolytic subunit
COX: Cytochrome c oxidase
Cpn10: Chaperonin 10 (10 kDa heat shock protein)
CPT1b: Carnitine palmitoyl transferase 1b
eIF2α: Eukaryotic translation-initiation factor-2 alpha
ER: Endoplasmic reticulum
ETC: Electron transport chain
FA: Fatty acids
FUNDC1: FUN14 domain containing 1
GAPDH: Glyceraldehyde-3-phosphate dehydrogenase
HAF-1: ABC (ATP Binding Cassette) transporter
H_2_DCF-DA: 2’,7’ dichlorodihydrofluorescein diacetate
HO-1: Heme oxygenase-1
HRP: Horseradish peroxidase
HSP: Heat shock protein
HSP60: 60 kDa heat shock protein
H2B: Histone 2B
IMF: Intermyofibrillar
IMM: Inner mitochondrial membrane
IMS: Intermembrane space
ISR: Integrated Stress Response
JNK: c-jun N-terminal kinase
KO: Knockout
kDa: Kilodaltons
LAMP1: Lysosomal-associated membrane protein 1
LC3: Microtubule-associated proteins 1A/1B light chain 3A
LDHA: Muscle-specific lactate dehydrogenase
LONP1: Lon protease 1
MnSOD: Manganese-dependent superoxide dismutase
mRNA: Messenger RNA
mtDNA: Mitochondrial DNA
mtHSP70: 75 kDA mitochondrial heat shock protein
mPTP: Mitochondrial permeability transition pore
NQO1: NAD(P)H quinone dehydrogenase 1
NR: Nicotinamide riboside
NuGEMPs: Nuclear genes encoding mitochondrial proteins
OMM: Outer mitochondrial membrane
OSNs: Olfactory sensory neurons
OXPHOS: Oxidative phosphorylation
PCR: Polymerase chain reaction
PFK-1: Phosphofructokinase-1
PGC-1α: Peroxisome proliferator activator receptor (PPAR) γ coactivator-1 alpha
PIM: Protein import machinery
PKR: Protein kinase RNA-activated
PQC: Protein quality control
RNA: Ribonucleic acid
ROS: Reactive oxygen species
SDS-PAGE: Sodium dodecyl sulfate polyacrylamide gel electrophoresis
SS: Subsarcolemmal
TFAM: Mitochondrial transcription factor A
uORF: Upstream open reading frame
UPR: Unfolded protein response
UPR^ER^: Endoplasmic reticulum UPR
UPR^mt^: Mitochondrial UPR
ATPase: Vacuolar-type ATPase
VDAC1: Voltage-dependent anion channel 1
WT: Wild-type

## Acknowledgements

We would like to thank Dr. Stavros Lomvardas at Columbia University for providing us with ATF5(+/-) heterozygous KO mice, in addition to Hani J. Shayya for assistance with breeding and genotyping. This work was supported by funding from the Canadian Institutes for Health Research (CIHR) and the Natural Sciences and Engineering Research Council of Canada (NSERC) to D.A. Hood. D.A. Hood is also the holder of a Canada Research Chair in Cell Physiology.

## Supplemental

**Table S1.**
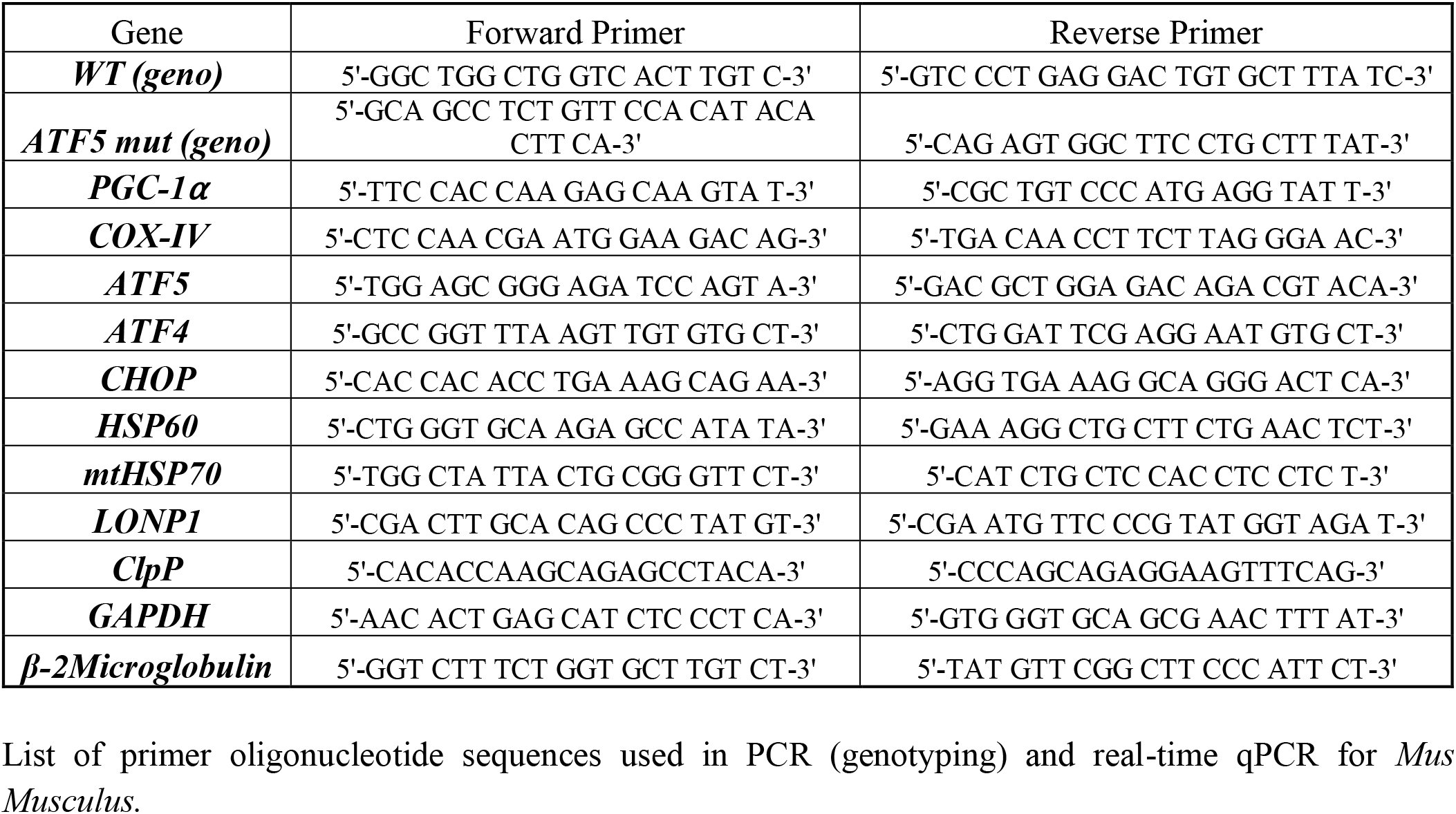
List of primer oligonucleotide sequences used in PCR (genotyping) and real-time qPCR for *Mus Musculus*.

**Table S2.** List of antibodies used in Western Blotting.

| **Antibody** | **Manufacturer** | **Catalogue #** |
| --- | --- | --- |
| **α-tubulin** | Calbiochem (Millipore) | CP06-100μg |
| **H2B** | Cell Signaling | 2934S |
| **GAPDH** | Abcam | ab8245 |
| **Aciculin** | Made in-house | N/A |
| **PFK-1** | Santa Cruz | sc-166722 |
| **LDHA** | Cell Signaling | 2012S |
| **CPT1b** | Proteintech | 22170-1-AP |
| **ATG7** | Sigma | A2856 |
| **Beclin-1** | Cell Signaling | 3738S |
| **LC3** | Cell Signaling | 4108S |
| **Parkin** | Cell Signaling | 4211S |
| **LAMP1** | Abcam | ab24170 |
| **V-ATPase** | Santa Cruz | sc-55544 |
| **NQO1** | Abcam | ab34173 |
| **HO-1** | Abcam | ab13248 |
| **MnSOD** | Upstate Cell Signaling Solutions | 06-984 |
| **Cytochrome c** | BD Biosciences | 556433 |
| **p-SAPK/JNK (T183)** | Cell Signaling | 4668S |
| **t-SAPK/JNK** | Cell Signaling | 9252T |
| **p-eIF2α (S51)** | Invitrogen | 44728G |
| **t-eIF2α** | Cell Signaling | 9722S |
| **ATF5** | Abcam | ab184923 |
| **ATF4/CREB2** | Santa Cruz Biotechnology | sc-390063 |
| **CHOP** | Cell Signaling | 2895S |
| **VDAC1/Porin** | Abcam | ab14734 |
| **PGC-1α** | EMD Millipore | AB3242 |
| **LONP1** | Cell Signaling | 28020S |
| **HSP60** | Enzo Life Sciences | ADI-SPA-806-D |
| **mtHSP70** | Enzo Life Sciences | ADI-SPS-825-F |
| **Cpn10** | Enzo Life Sciences | ADI-SPA-110-D |
| **Anti-Mouse HRP-linked 2°** | Cell Signaling | 7076S |
| **Anti-Rabbit HRP-linked 2°** | Cell Signaling | 7074S |

